# The vigor paradox: saccade velocity during deliberation encodes utility of effortful actions

**DOI:** 10.1101/2022.03.09.483677

**Authors:** Colin C. Korbisch, Daniel R. Apuan, Reza Shadmehr, Alaa A. Ahmed

## Abstract

During deliberation, as the brain considers its options, the neural activity representing the goodness of each option rises toward a threshold, and the choice is often dictated by the option for which the rise is fastest. Here we report a surprising correlate of these activities: saccade vigor. We engaged human subjects in a decision-making task in which they considered effortful options, each requiring walking various durations and inclines. As they deliberated, they made saccades between the symbolic representations of those options. These saccades had no bearing on the effort that they would later expend, yet as they deliberated, saccade velocities increased. The rate of rise in vigor was faster for saccades toward the option that they later indicated as their choice, and encoded the difference in the subjective value of the two effortful options. Once deliberation ended, following a brief delay the subjects indicated their choice by making another saccade. Remarkably, vigor for this saccade dropped to baseline and no longer encoded subjective value. These results are consistent with an urgency model of decision-making in which a global signal in the brain drives both the neural circuits that make decisions, and the neural circuits that make movements. Paradoxically, this common drive is shared between the oculomotor circuits and the decision-making circuits, even when the decision involves effortful expenditure during a future event.

**Significance:** There is a link between the decisions we make and the movements that follow. Not only do we prefer options of greater value, but we also move faster to acquire them. When deliberating between options, neural activity rises to a threshold and the option that wins this race is the one chosen. We report a potential correlate of this in the motor control circuits; during deliberation, saccade vigor to both options rise, but faster for the option ultimately chosen. Thus, our movements appear to mirror the neural activity conducting the decision-making process. Paradoxically, this is true even when the movements have no direct bearing on the decision at hand.

## Introduction

Imagine an autonomous humanoid robot that employs its head-mounted cameras to gather information from its environment, relies on its decision-making algorithm to assign utility to each available option, and then moves its limbs to act on the option with the greatest utility. The choice that this robot makes, and even the algorithm, might be indistinguishable from a biological brain that consumes the same visual information. However, in one respect the robot’s actions differ: our brain moves our eyes with a vigor that depends on the utilities of each option, but not the robot. For example, as humans and other primates deliberate between rewarding options, their saccadic eye movements exhibit peak velocities that reflect the subjective value of the expected reward. This suggests that in our brain, the neural circuits that make decisions are inexorably linked to the neural circuits that make movements. However, therein lies a paradox.

To explain this paradox, consider that from a theoretical perspective, the propensity for reward to invigorate movements is consistent with an optimization principle (1, 2). In this framework, animals make decisions and movements that maximize the sum total of costs and rewards, divided by time (3). Moving faster to acquire reward is justified because the increased effort expended in making a faster movement is offset by acquisition of an earlier reward, which increases the rate of reward (4) and improves fitness (5). However, the paradox lies in the fact that if we reach faster, or walk faster, we will acquire the reward earlier, but faster saccades will not hasten the reward.

For example, as we consider two rewarding options presented on a video monitor, we direct our gaze from one image to the other. Initially, the velocities of saccades directed toward the images are comparable (6, 7), but as deliberation proceeds, saccade velocity toward one image becomes larger, foretelling the choice. This observation suggests that the vigor of a saccade toward a rewarding object reflects the value that the brain has assigned to that object. But why should this be?

One possibility is that there is a common signal that is shared between the decision-making circuits and the motor circuits. A candidate for this signal is urgency, a signal that during deliberation gradually increases, pushing the decision-making circuits towards a threshold that determines the choice (8). In this model, the immediate value for each option is weighted by the urgency signal. As a result, the option that has a greater value rises faster, reaches the threshold earlier, and is thus chosen. Correlates of this signal have been noted in the basal ganglia (9), a structure that directly influences saccade vigor (10, 11).

If urgency is a signal that links the decision-making circuits with the saccade circuits, then we should see several behavioral consequences. During deliberation, as urgency monotonically increases, so should saccade vigor, but the rate of rise in vigor should be faster for saccades that are directed toward the more valuable option. Moreover, once the deliberation period is over and the urgency signal returns to baseline, so should saccade vigor.

To evaluate these predictions, we considered a decision-making task that involved effortful options. People deliberated between options that involved walking across various inclines and distances. They naturally made saccades between the symbolic representations of these options, but the saccades had no bearing on the effort that they would expend in the future. Importantly, we separated the deliberation period from the choice period, thus separating saccades that were made during a theoretical rise in the urgency signal, and those that were made when urgency has returned to baseline. Would saccade vigor reflect subjective value even in this case in which the cost of effort was paid by effectors that were completely distinct from the oculomotor systems? Once the deliberation period was over, would saccade vigor return to baseline, even though that saccade indicated choice?

If saccade vigor reflects subjective value, then a saccade towards the less costly option should exhibit a greater vigor. However, saccades are also influenced by the salience or the attentional demands of the option (12): an option that carries a greater cost might elicit greater saccade vigor because of greater aversion. For example, activities of neurons in the parietal cortex increase with the reward value of the option, but are also greater if the option carries a greater cost (13). Thus, this view would predict that saccade vigor will be greater when the movements are directed toward more effortful options because those options carry greater attentional demands.

It is also possible that saccade vigor during decision-making would show no relationship to effort costs. Manohar et. al. (14) recorded saccade velocities in response to either a behaviorally contingent (in that the participants could influence the outcome) or random penalty. They found that if the loss was contingent on reaction time, saccade peak velocity was larger for the greater expected loss. However, on the random trials in which the loss amount was guaranteed, varying the expected loss produced no changes in saccade vigor. This view suggests that if the decision is between two effortful options, saccade vigor will not be affected by the expected effort because that effort is not contingent on the vigor of the saccade.

Contrary to both predictions, but consistent with the urgency model, here we found that saccade vigor increased during deliberation, with a rate that was faster when the movements were directed towards the less costly option. When the deliberation period ended, participants indicated their choice by making another saccade. Vigor dropped following the completion of the deliberation period, and no longer reflected the effort cost of the chosen option.

## Results

In our study, participants considered walking various inclines and durations. Their options were represented by a visual symbol on a computer monitor. We tracked their eye movements as they deliberated and ultimately chose one of the two options. The decision-making process was divided into two parts. In part 1, participants deliberated and naturally made saccades between the options without any explicit instructions. When they arrived at a decision, they signaled by hitting a computer key, fixated at a center target, and then indicated their choice by making a saccade towards their preferred option (part 2). To emphasize that their choices had real consequences, we selected a random subset of trials and asked the participants to walk the chosen inclinations and durations.

The experiment began with a familiarization phase in which the participants (n=22) experienced various inclines by walking on a treadmill and learned to associate each with a graphical symbol (Fig. 1A). Following familiarization, they deliberated between the options (Fig. 1B), which differed along two dimensions: walk incline and walk duration. Walk speed was fixed at 3.1 m/s. One of the options was always the reference: 4% incline at 6 min duration, or 5% incline at 5.5 min duration. Participants had up to 4.5 sec to arrive at their decision. They indicated this by pressing the spacebar, after which a fixation cross was presented on the screen. Following a brief delay (1.5±0.25 sec, mean ± s.d.), they indicated their choice by making a saccade towards a grey target on the same side of the screen previously occupied by the option they preferred (Fig. 1B).

**Figure 1.**
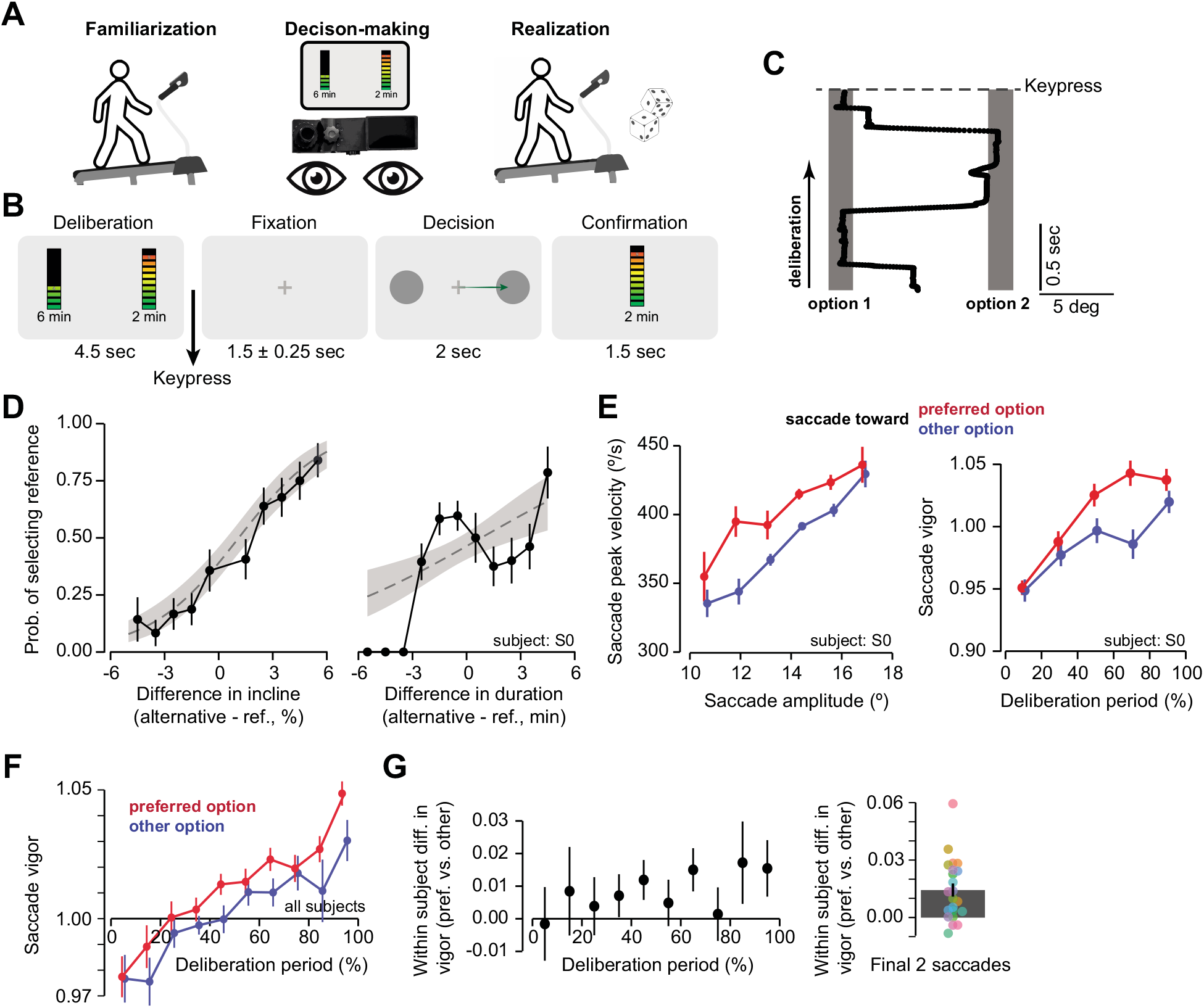
A) Outline of experimental protocol. The experiment was comprised of three steps: familiarization, which involved repeated exposure to walking at various inclines on a treadmill; decision-making, where participants’ eye movements were recorded as choices were made; and realization, when a random subset of previous choices were realized on the treadmill. B) Effortful choice selection was split into distinct phases: deliberation, fixation, decision, and confirmation. Participants were given limited amounts of time to deliberate their movement choices (maximum of 4.5 s) as well as making a saccade to confirm their movement decision (2 s). C) Example trajectory of eye movements over the course of deliberation. The two graphics representing the movement options were placed at approximately 1/3rd and 2/3rds of the horizontal distance of the computer monitor, a 14-degree visual angle separation on average across participants. Once a choice was in mind, participants could conclude the deliberation phase before the maximum duration of 4.5 seconds by pressing the keyboard’s spacebar. Each dot represents a gaze localization sampled at 1000 Hz. D) Example participant S0 demonstrated sensitivity in their choices to both incline and duration differences. Grey regions represented 95% CI for logistic fit. E) Saccade vigor profiles for example participant. *Left* Sub-selection of saccades which horizontally traversed the computer monitor (see Figure 1C). Peak velocity was greater towards the preferred option. *Right* Participant’s saccade vigor was greater towards the preferred option as compared to the nonpreferred option as deliberation progressed. F) Across all participants, saccade vigor increased over the course of deliberation and was significantly greater towards the preferred option at the end of deliberation. G) *Left* The difference in vigor between the two options increased over the course of deliberation. *Right* Highlighting the difference in vigor for only the final two saccades. Individual participant vigor differences represented as colored points.

### Saccade vigor increased during deliberation

During a typical trial, the subjects made saccades from one option to the other, as shown for a representative subject in Fig. 1C. This subject had a propensity towards choosing options with lower relative inclines and durations (Fig. 1D), as expected for a desire to minimize energetic costs. Intriguingly, during the deliberation period the saccades toward each option had velocities that reflected this preference. Peak saccade velocity as a function of amplitude was greater when it was directed towards the option that was ultimately chosen (Fig. 1E, left panel). To quantify this effect, following our earlier work (6, 15) we defined vigor of a saccade as the ratio between the actual peak velocity and the participant’s expected peak velocity for a given amplitude. The expected peak velocity was defined as:

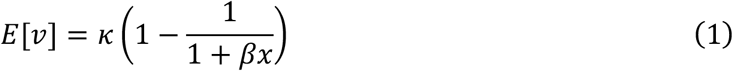

In this expression, *κ* and β were parameters specific to each subject, *x* was saccade amplitude, and *E*[*v*] was the expected value of peak velocity. Vigor of saccade *n* was defined as the ratio 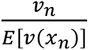. Eq. (1) was fitted to the saccades of each participant and then used to determine the vigor of each saccade. For this participant, vigor near the end of the deliberation period appeared larger if the movement was directed towards the ultimately chosen option (preferred) (Fig. 1E, right panel).

When we considered the data across all participants, we found that as they deliberated, saccade vigor gradually increased by 4.93±1.013% (LMM, p=0.00107). The rate of increase was faster for saccades directed towards the preferred option (β=6.726e-3 ± 6.66e-3 [±95% C.I.], p=0.0478) (Fig. 1F). When we considered the entire deliberation period, saccade vigor towards the preferred option was significantly greater (β=1.29e-2±5.29e-3, p=1.84e-6) (Fig. 1G, left panel). This effect of preference on vigor was particularly prominent in the final two saccades of the deliberation period (t_(21)_=-3.165 p=0.00467; Figure 1G, right panel).

The rise in saccade vigor during the deliberation period resembled theoretical variables that are commonly employed in models of decision-making. According to these models, as the subject deliberates there is a race between variables that reflect the goodness of each option. For example, when the decision is easy and thus the deliberation period is brief, the decision-variable toward the preferred option rises faster than when the decision is hard. How did saccade vigor evolve during brief or long deliberations?

For each subject we computed their median duration of deliberation and normalized each deliberation period with respect to that value. We found that saccade vigor increased more rapidly when the deliberation period was short (Fig. 2A, left subplot, time × short/long interaction; β=-1.95e-2±8.66e-3, p=1.03e-5). Remarkably, the faster rate of increase in vigor was present in the short deliberation trials even when we considered only those saccades that were made toward the reference option (Fig. 2A, right subplot; β=-2.33e-2±1.18e-2, p=1.2e-4). Similarly, in short deliberation trials the saccades that were made toward the preferred option exhibited a rise in vigor that was faster than compared to long deliberation trials (Fig. 2B, left subplot; β=-2.19e-2±7.43e-3, p=7.6e-9).

**Figure 2.**
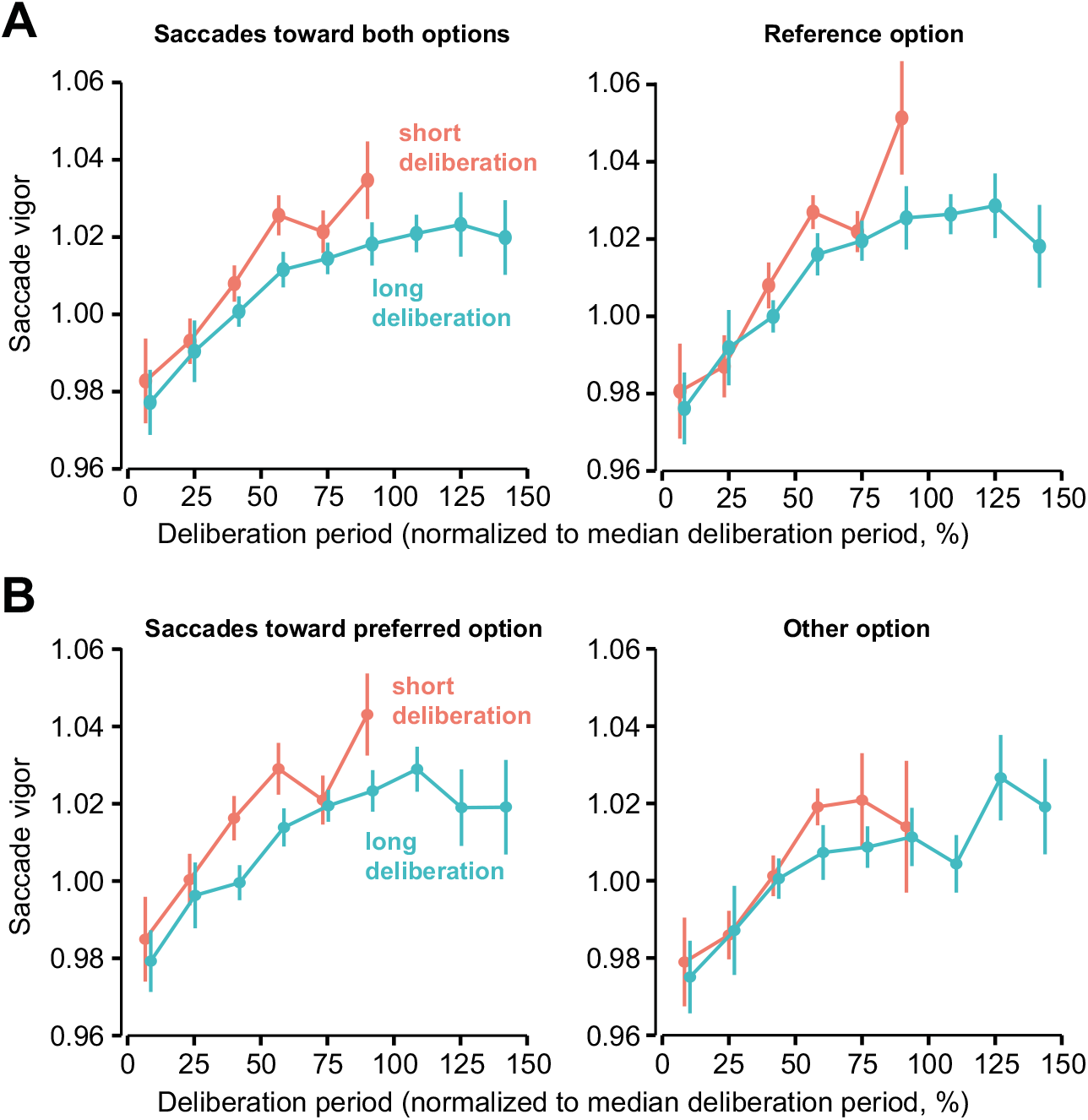
A) Participants’ rate of change in vigor over the course of deliberation differed between short and long deliberations. Short deliberations were defined as trials shorter than participant median time, and long deliberations were greater than this time. Saccade vigor data are plotted against time normalized as percentage of participant median deliberation. *Left* Data averaged across all saccades. *Right* Data averaged only for saccades directed towards the reference option. Even though the absolute value did not change, rate of vigor increase when directed towards the reference significantly differed between short and long deliberations. B) Data are presented separated by preference (whether directed towards option ultimately chosen or no). Rate of rise in vigor for preferred saccades also differed between short and long.

In summary, saccade vigor increased during deliberation. The rate of rise was faster for saccades that were directed toward the preferred option and was faster in trials in which the deliberation period was shorter.

### Saccade vigor indicated degree of preference

Is saccade vigor a binary measure of preference, or a continuous indicator of degree of preference? To answer this question, we tried to quantify the subjective cost, or the utility, that each subject assigned to each option. This required fitting a model to the choices that the subjects had made.

We began by checking whether the choices were rational: did the subjects prefer the option that was less effortful? Indeed, as the difference between the incline of the alternative option increased with respect to the reference, the participants were more likely to choose the option with the lower incline (GLMM, β=0.713±0.0879, p<2e-16, Fig. 3A). Similarly, the participants were more likely to choose the option with lower duration (β=1.38±0.096, p<2e-16, Fig. 3A). Next, in order to quantify the subjective cost that each participant assigned to an option, we calculated the expected metabolic cost (16, 17) in units of Joules for a person of mass *m* to walk a duration *t* on an incline *g* with average velocity *v*:

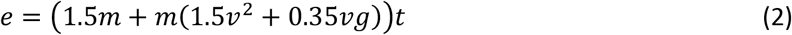

**Figure 3.**
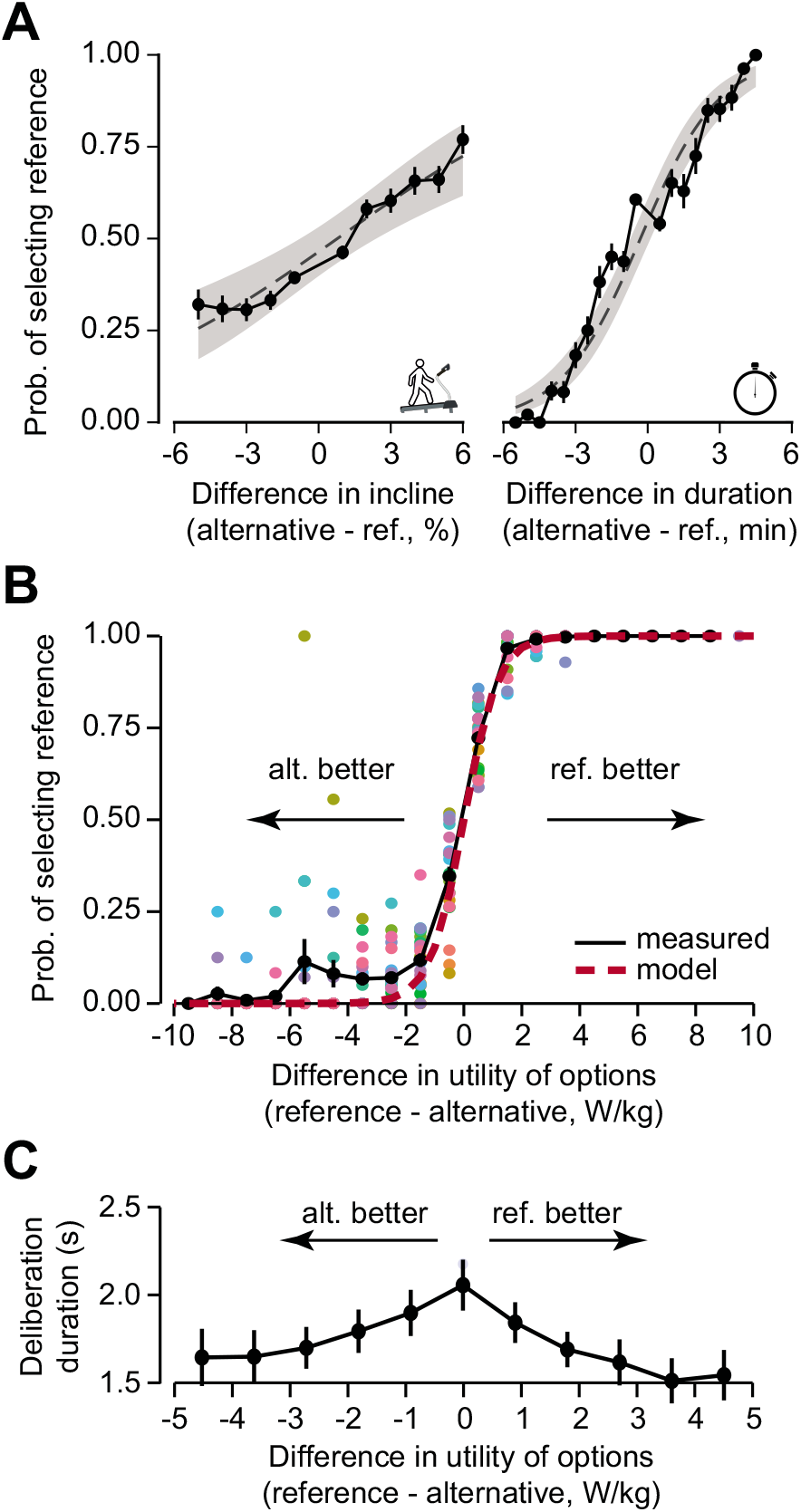
A) Participants showed consistent sensitivities to incline and duration. Shaded region represents 95% CI of logistic fit. B) Utility difference provided a more accurate predictive model for participant choices than incline and duration combined. Dotted line represents logistic regression fit. C) Deliberation time varied with capture rate difference. Across participants, average deliberation time was longer for smaller capture rate differences.

The above expression represented the expected energetic cost of the walk. In this equation, all parameters are known and thus there is nothing to fit. Next, we estimated the utility of an option via the difference between the expected reward and energetic cost, divided by duration of the walk:

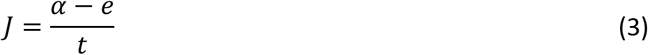

The above expression represents the capture rate (1). In Eq. (3), the only unknown parameter is *α*, the subjective value of reward, which we estimated by fitting the equation to the choices that the participants had made (see Methods). The result was a utility, or goodness, of each option. The capture rate correctly predicted 86.6±1.2% (mean ±s.e.) of choices that the participants made (Fig. 3B). However, the model had limitations, as it could not account for the fact that the participants had a greater preference than expected for the reference option (Fig. 3B, region of the data in which difference in utility of options is negative).

We compared this model with an alternative one in which each subject had a specific sensitivity to incline and duration. In this alternative model, the probability of selecting the reference option was a function of the difference in incline and difference in duration between the reference and the alternative (see Methods). Comparing the two models, we found the capture rate model (Eq. 3) provided a more parsimonious explanation for choice behavior (AICs: 3560.8, 4430.1 respectively).

Equipped with this quantitative description of utility for each option (Eq. 3), we tested it with the behavioral metric of deliberation time (18–20). Deliberation time is expected to increase for options with a smaller difference in capture rate (i.e., a difficult decision). As hypothesized, we found that deliberation time varied robustly with participants’ capture rate difference (GLMM, β = −6.032±2.116, p=2.4e-8; Fig. 3C). Deliberation was longest for capture rate differences near 0 J/kg·s and decreased as options differed in value; thus, the greater the difference in the capture rate of the two options, the more quickly participants were able to make their decision. Capture rate provided us with a participant-specific measure of the utility of each option, and the difference between the utility of each option provided a measure of the degree of preference. Armed with these quantities, we returned to our original question: does saccade vigor signal which option is preferred, or does it also signal how much one option is preferred over the other?

We examined saccade vigor on easier trials compared to more difficult ones to determine whether the saccades revealed degree of preference. Looking again at our example participant, we separated trials with the greatest 20% and smallest 20% differences in capture rate, thus identifying the easiest and most difficult trials. In easy trials, saccade vigor appeared to increase at a faster rate than in the hard trials (Fig. 4A). Across all participants, saccade vigor increased during deliberation (main effect; β=2.026e-2±7.434e-3, p=3.43e-5). However, the rate of increase in vigor (the slope of the relationship with time), was influenced by the capture rate differential of the two options (interaction term **|rate| × time**; β=3.281e-4±2.38e-4, p=0.00688). Thus, the greater the difference in the utility of the two options, i.e., the easier the decision, the greater the change in saccade vigor during deliberation.

**Figure 4.**
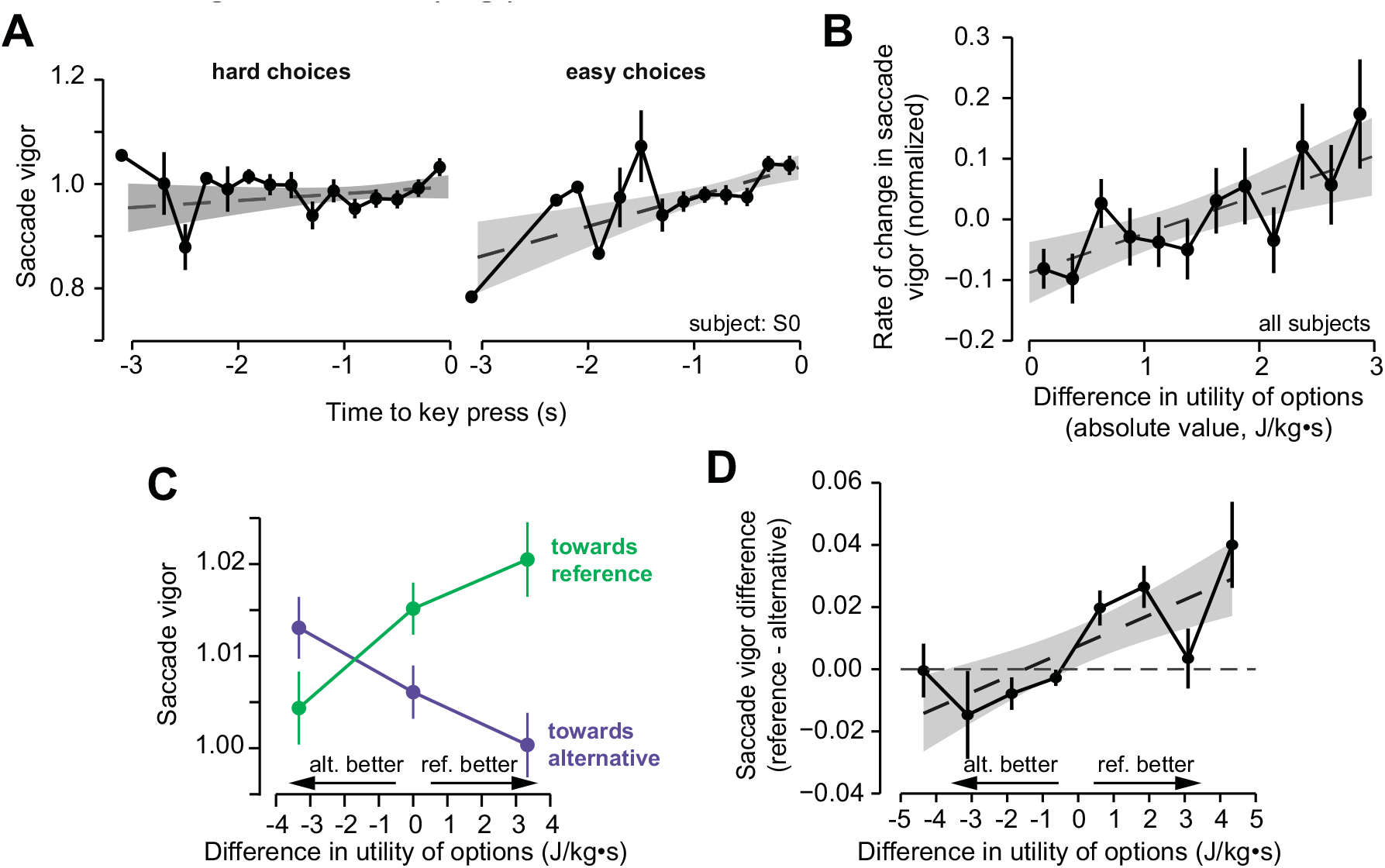
A) Subject 0 showed difference in vigor rate increase during deliberation dependent on capture rate difference. *Left* For trials with the smallest capture rate differences (i.e. more difficult choices), vigor rate was relatively flat. *Right* Those trials with greater differences in capture rate had steeper rates of vigor increase up to the end of deliberation. B) For all participants, the rate of saccade vigor increase (1/s) over the course of deliberation significantly increased with absolute capture rate difference. C) Saccade vigor towards reference and alternative options significantly varied with capture rate difference. As either option increases in relative utility, vigor towards that option increases. D) The difference in vigor towards the reference option and alternative option increased with capture rate difference. Shaded regions represent 95% CI.

To visualize these results, in each trial with at least two saccades, we calculated the rate of increase in saccade vigor (ΔVigor/ΔTime) and calculated its z-score per participant (Fig. 4B). We tested the correlation between this per-trial measure to the capture rate difference and again, we found the effect to be significant (p=0.0378). We then asked whether target preference modified the rate of vigor change and found that the rate of change varied with absolute capture rate difference for saccades directed towards the preferred option (**|rate| × time × preference**; β=1.361e-3 ± 1.159e-3, p=0.0214). Thus, the greater the degree of preference, the greater the rate at which saccade vigor increased over the deliberation period.

We next considered saccades towards the reference or alternative options, rather than towards the preferred or nonpreferred options. This has the effect of making changes in relative utility linear. When categorizing the final two saccades as either directed towards reference or alternative, we found the average vigor of saccades towards the reference increased with increasing relative capture rate (β=4.785e-3±2.017e-3, p=3.36e-6; Fig. 4C). Saccade vigor towards the alternative option decreased with decreasing relative utility (β=-2.38e-3±1.435, p=0.0011). Thus, when one option had a larger capture rate than the other, saccade vigor towards that option was also greater.

We further examined whether the difference in saccade vigor between the reference and alternative options reflected the underlying difference in their utilities, i.e., whether the difference in vigor reflected an individual’s degree of preference for one option over the other. We calculated the per-participant difference in average vigor between saccades directed towards the reference and the alternative if made as part of the final two saccades per trial, binned by capture rate difference (bin count = 8, 1298±871 saccades per bin, mean±s.d.). We then measured the correlation between these mean differences and utility. We found a significant relationship between the relative vigor and relative utility (ρ=0.774, p=0.02422; Fig. 4D).

In summary, saccade vigor increased as subjects deliberated between two effortful options. As the choice became easier, vigor increased at a faster rate. Near the end of the deliberation period, the differential in vigor for saccades directed toward the preferred option versus the other option was greater when the difference in the utilities of the two options was greater. Thus, the rate of increase in saccade vigor and relative vigor of the final saccades appeared to reveal the degree of underlying preference.

### Reaction time of the decision saccade, but not its vigor, varied with preference

Our experiment separated the deliberation period from the period in which the subject revealed their choice. In both periods the subjects made saccades. However, in the deliberation period the saccades were made for the purpose of acquiring information, whereas in the final phase the saccade indicated choice. Did the choice-period saccade carry information regarding utility, or was the link between utility and vigor present only when the oculomotor system was actively participating in the acquisition of information?

Once the deliberation period was over (press of a key), the participants fixated upon a centrally located stimulus and, after a variable delay, two grey circles appeared on either side of the screen, corresponding with the previous two options. The appearance of the circles cued the participants to make a saccade and indicate their choice. The time it took to initiate this saccade, i.e., reaction time, was correlated with the capture rate difference between the options (β=-1.054±0.657, p=0.00168). The greater the capture rate difference, the shorter the reaction time (Fig 5A).

**Figure 5.**
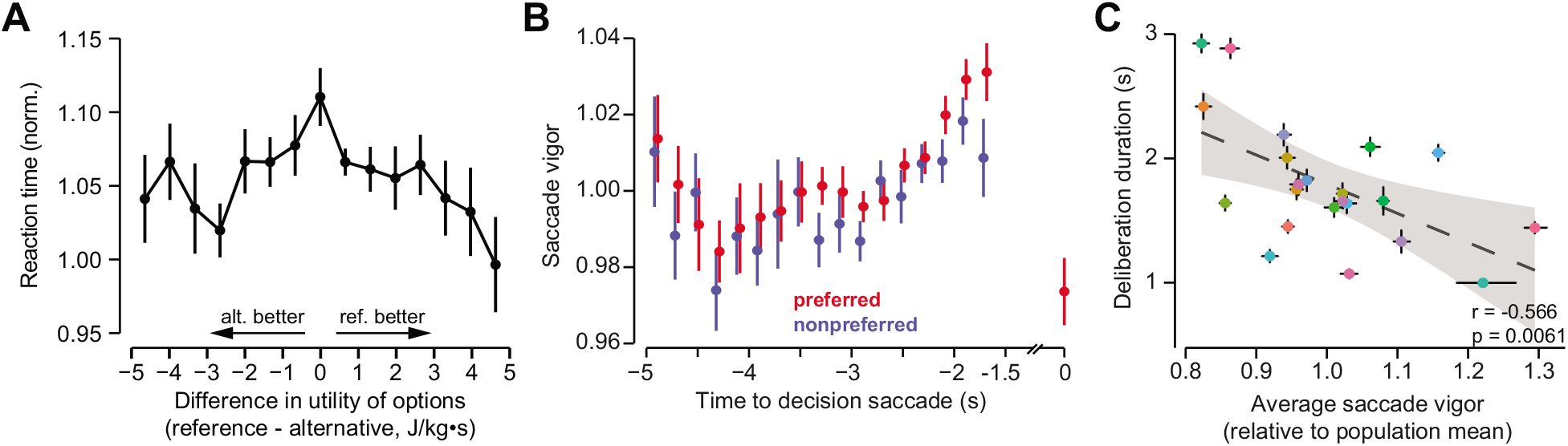
A) Reaction time was found to significantly vary with capture rate difference. With increasing capture rate difference, reaction times decreased (p=0.002). Participants’ reaction times were normalized to median values. B) Decision saccade vigor was significantly less than the preceding final saccade in the deliberation phase (p=0.0002). C) Significant correlation was found between participant saccade vigor and their average deliberation time. Participants who deliberated more quickly made relatively faster saccades. Shaded region represents 95% CI.

However, there was a significant drop in saccade vigor from the end of the deliberation phase to the decision phase (β = 0.046±0.02, p=0.000162; Fig 5B), which increased over the course of the experiment (GLMM, **phase × trial** interaction, β = −0.03399±0.012, p=6.45e-8). Furthermore, unlike the saccades at the end of the deliberation period, the vigor of the decision saccade did not vary with capture rate difference. Measured against this difference, the fitted coefficient was found to be indistinguishable from zero (β=-0.0699± 0.223, p=0.538). Even when separately considering the selection of the reference or alternative option, there was still no effect (β=2.428e-5±8.848e-5, p=0.784).

Thus, saccade vigor rose during deliberation but dropped back to near baseline when deliberation ended. Saccade vigor during deliberation was sensitive to the difference in the utilities of the two options, but not when the saccade signaled the choice. The easier the decision, the shorter the reaction-time of this decision saccade.

### Individuals who had greater saccade vigor tended to have shorter deliberation periods

Some individuals persistently make saccades with high velocities, while others have saccades that are much slower (15, 21–23). These vigor differences are not a reflection of a speed-accuracy tradeoff, as people who make faster saccades are not sacrificing endpoint accuracy for the purpose of arriving sooner (15). Thus, saccade vigor may be a trait that dissociates people. Is there a relationship between how an individual evaluates reward, effort, and passage of time for the purpose of decision-making, and how they evaluate these same variables for the purpose of making a movement?

A meta-analysis of saccade velocities in healthy people and those with neurological disorders suggested that groups that tended to make impulsive decisions also exhibited increased saccade velocities (24). Two recent reports by Berret and colleagues suggest that in the context of arm movements, vigor differences among healthy people reflects a propensity to expend effort in order to save time (25, 26). That is, cost of time tends to be greater in people who move with greater vigor. Our experiment allowed us to ask whether there was a relationship between an individual’s saccade vigor, and their decision-making patterns.

To compare saccade vigor across participants, we fit a hyperbolic function for all saccades, producing a population-averaged expected velocity as a function of amplitude (15). We next defined the vigor of each saccade for each subject with respect to this expected value, and then averaged the vigor of all saccades for each subject. The result was the participant’s average saccade vigor with respect to the population mean.

We found that those participants who had greater than average saccade vigor allocated significantly less time for deliberation (t_(20)_=-3.068, ρ=-0.566, p=0.00607; Fig. 5C). In other words, participants who tended to make faster saccades also tended to deliberate for a shorter period. This difference in vigor could not account for the range of deliberation times measured over the experiment, as the maximum per-saccade difference in vigor (from the slowest to the fastest) amounted to approximately 15 milliseconds whereas the range in average deliberation times was near 2 seconds.

### Control studies

The low-level properties of a visual stimulus can bias the decisions that subjects make. For example, when utilities of the stimuli are similar, people are biased toward choosing the brighter stimulus (27). During the deliberation period, the symbols that represented the effortful options had differing luminosity. Salience is greater for stimuli that have greater luminosity, thus raising the concern that differences in vigor may be related to luminosity. However, we designed our task so that luminosity was anti-correlated with capture rate (t_6606_=17.9, p<2.2e-16). That is, the more effortful options were represented by more luminous symbols. Yet, saccade vigor was greater toward the less effortful options. Beyond that association, saccade vigor was not affected (GLMM, β=-1.259e-2±2.04e-2, p=0.227).

## Discussion

As people deliberated between two options that promised future effort, their saccades increased in vigor, but the rate of rise was faster for saccades that were directed toward the less effortful option. Indeed, the difference in the rates of rise in vigor reflected the utility differential. Once the deliberation period ended, even though the choice was signaled via a saccade, vigor returned to baseline and no longer conveyed the utility of the chosen option. Remarkably, people who had higher saccade vigor with respect to our population mean tended to deliberate for a shorter period, arriving at their decision earlier than those who had low saccade vigor.

These results suggest a link between the neural mechanisms that control movement vigor, and those that evaluate the effort cost of an option. Previous work has established that dopamine release plays a causal role in invigorating movements (28), yet in forced-choice scenarios, dopamine is poorly modulated by effort costs (29). The fact that saccade vigor reflected effort costs during deliberation but not choice readout, predicts that if dopamine is the principal modulator of vigor, then it likely encodes effort costs during deliberation.

### Effort cost of the option modulates saccade vigor during deliberation

From a normative perspective, greater potential for reward should invigorate movements because faster movements hasten reward acquisition, which in turn increase the rate of reward (1). However, when deliberating between options, there is no theoretical rationale for changes in saccade vigor because these movements have no significant bearing on when the chosen option will be exercised. Thus, changes in saccade vigor during deliberation cannot alter the rate of reward or effort. Yet, during deliberation, Thura and Cisek found that saccade vigor increased while monkeys deliberated between two potentially rewarding options (30). If saccade vigor reflects a utility signal common to both decision making and movement control, then vigor should be influenced by effort costs of the options as well. Here, we found that saccade vigor increased during deliberation between two effortful options, but at a faster rate for the option that had the lower effort cost. Remarkably, saccade vigor returned to baseline once the deliberation period ended, even though this saccade was critical for signaling choice.

Saccade vigor is closely linked to the activities of neurons in the superior colliculus (31), which in turn receive information regarding the utility of the visual stimulus from the frontal eye field, parietal cortex, and the basal ganglia (1). The information that is conveyed regarding the value of the visual stimulus to the superior colliculus is a balance of excitation from the cerebral cortex, and inhibition from the basal ganglia. When the stimulus has a greater reward value, the inhibition that substantia nigra reticulata (SNr) neurons impose on the superior colliculus is suppressed, which encourages a high vigor saccade toward that stimulus (32). SNr activity reflects utility calculations in the caudate (33), which are conveyed to the SNr via a direct pathway that reflects reward value of the stimulus, and an indirect pathway that reflects the risk or cost of that stimulus (34). The fact that saccade vigor was greater toward the less effortful option makes a critical prediction: during deliberation, activities of the indirect pathway neurons in the caudate will be greater when the saccade is directed toward the more effortful option (35).

Why does saccade vigor reflect effort cost of the various options during deliberation? One possibility is that the decision-making and motor control circuits of the brain are both modulated by an evolutionarily ancient neurotransmitter, dopamine. Release of dopamine near movement onset invigorates many kinds of movements, including saccades, reaching and walking (36–39). Expectation of reward promotes dopamine release in a wide range of capacities. It is possible that dopaminergic circuits not only signal the utility of the option, but also influence movements across modalities, even when the movement does not affect reward rate.

However, if dopamine release is the basis of increased saccade vigor during deliberation, then the fact that our results were in the context of effortful choices would imply that dopamine release is greater in response to stimuli that are associated with lower effort. This appears inconsistent with the available data: in a forced-choice task, dopamine neurons responded to the reward value of the stimulus, but not its effort cost (40, 41). In contrast, during deliberation the dopamine levels in the ventral striatum are greater when a given reward can be acquired with low effort as compared to high effort (42). Indeed, blocking D1 dopamine receptors in the ventral striatum reduces the animal’s willingness to exert effort for a given reward (43).

Our results provide a potential solution to this puzzle by noting that saccade vigor reflected effort costs, but only during deliberation. Our design contrasted with most previous studies in which the act of the saccade was itself associated with a reward or punishment (7, 44–46). In our design the time of the decision and the time of its readout were separated by at least three seconds. Saccade vigor dropped after the time of the decision and no longer reflected the options’ relative value. If vigor is a proxy for dopamine release (28, 39), then our behavioral results predict that during deliberation, there will be greater release before saccades that are directed toward the less effortful option. Thus, deliberation may be the key period during which there is a link between the decision-making and the motor control circuits.

Outside of a motor control context, researchers have sought to investigate the link between dopamine and avoidance of aversive outcomes (47–49). Avoidance of aversive or more effortful outcomes is rewarding in and of itself (50). Thus, reinforced behavior from active avoidance may also be modulated by dopamine (51). As of now, there is no definite evidence for a link between negative reinforcement and dopamine in humans (52).

Dopamine-modulated or not, there is evidence that vigor of movements depends on the expected effort cost of that movement. Reaching movements are slower when moving greater mass and over greater distances (4, 15). Saccades are slower to more eccentric targets, possibly due to the greater effort needed to maintain gaze at increased eccentricities (2). Interestingly, this slowing is not necessarily specific to the effector exerting the greater effort. Individuals who move their head faster also tend to reach faster, and those who reach faster tend to walk faster (53). It is possible that effort-based net utility is represented beyond dopaminergic circuits, and that this representation influences vigor across movement modalities.

### Vigor during deliberation is dependent on utility, not salience

Previous studies concerning how saccade vigor related to a rewarding outcome were unable to fully distinguish between the value itself and the salient representation of the saccade target (6). It is unclear whether participants responded to the value or merely its perception, as a greater positive value might provoke greater salience/prominence. When both targets represent positive utility outcomes (e.g., individuals are rewarded with juice or money) then salient representation is indistinguishable to the reward value: greater reward suggests greater salient representation.

Salient representation may have more to do with motivation to avoid a bad outcome rather than a potential gain, as exemplified by loss aversion. Indeed, many neurons in the parietal cortex show greater activity for stimuli that promise greater reward, and stimuli that promise greater punishment (13). This might predict that saccade vigor will be greatest when directed towards the more effortful option. However, we found the opposite result; vigor was greater for saccades directed towards the less costly option. Kobayashi et al. also found supporting evidence of such a relationship when investigating motivational salience and saccade vigor (54). When training macaques on a visually guided saccade task, saccade vigor was greatest when a successful trial was rewarded, and least vigorous when avoiding a punishment.

### People who made faster saccades tended to make faster decisions

We found that individuals who tended to make saccades with greater vigor also tended to make decisions faster. One possible explanation lies in the dual effects of reward rate on vigor and decision making. A greater reward rate should lead to increased vigor. Deliberation between two options is commonly modeled as a choice between competing options, with evidence accumulating over time until a threshold is reached and the option is selected (55). In this framework, if evidence accumulation is a measure of reward rate, then a greater reward rate will also lead to an earlier threshold crossing and a faster decision. Thus, a trait-like subjective valuation of reward rate can provide an explanation for between-participant changes in vigor and decision making. This builds on previous findings where individuals who made faster saccades were also less willing to wait for reward (22), and those who reached with greater vigor tended to exhibit a greater cost of time (26). Berret and colleagues have suggested that people who exhibit greater vigor tend to have a greater cost of time (26). In a locomotion study, we found that individuals who preferred to run to a target (rather than walk), also tended to run with greater vigor for any distance (56). This potential link between individual differences in decision-making and movement control remains a fascinating area of further study.

### Vigor reflected relative utility with a potential exposure bias

Our subjects displayed a slight choice bias for the reference option, even when utility was equal (Fig. 3B). This bias toward the reference option was also present in saccade vigor. Vigor of the final deliberation saccade towards the reference option, regardless of time or utility, was significantly greater as compared to the saccade directed towards the alternative option (Fig 4C). This reference-bias may be accounted for by the exposure-dependent effects on subjective value assignment. Several studies have shown a relationship between increased gaze-duration and propensity for selection (57–61). Here, participants were exposed to one of the two reference images on every trial, i.e., 50% of trials, and so they experienced a much greater exposure to the reference options. Additionally, Krajbich et al. proposed that attention, measured as fixation duration, biases choice (62). Indeed, within a trial, we observed that participants tended to gaze for longer durations and were more likely to look last at the reference option. Thus, the greater exposure coupled with the greater gaze duration within a trial may have contributed to the greater value for the reference options.

### Limitations

The interval period between the deliberation and decision phase may have influenced decision-making and later saccade vigor. This period could have been used to reevaluate what decision was in mind given by the infrequent occurrence of high-vigor, last-look saccades directed towards the nonpreferred option. Although stated that the participant should know what they will later choose at this point, the potential for answer-switching is impossible to eliminate in this protocol.

### Conclusions

When considering prospective effortful walking options, the vigor of eye movements increased during deliberation, but at a greater rate toward the option that had the greater utility (less effortful option). The difference in the rate of rise in vigor reflected the degree of preference. Intriguingly, this relationship disappeared when the deliberation period end, and the saccade indicated the choice. Finally, people who had faster saccades also made faster decisions, implying that saccade vigor may also be predictive of decision-making behavior at a trait-like level.

## Materials and Methods

### Participants

Participants were recruited from the University of Colorado Boulder (n=22; 8 Female; age: 23.71±3.07, mean ± s.d.) with no known neurological or motor deficits. Each participant signed a consent form approved by the CU Boulder Internal Review Board and was paid $10/hr for participation in the study. Two subjects wore necessary corrective eyeglasses.

### Experimental design

Participants performed a decision-making task in which they considered walking on various inclinations and durations, each represented by a visual symbol on a computer monitor. A total of 360 decisions were presented per participant, with 180 unique decisions repeated twice, counterbalanced by left-right presentation order on-screen.

Participants were first familiarized to the various potential inclines to be decided between before any choices were presented. Half of subjects were familiarized to inclines of 0,2,4,6,8, and 10%, while the other half experienced inclines of 0,1,3,5,7, and 9% each for 3 minutes apiece. A brief reset period of 30s was allowed between each exposure. Familiarization order was randomized per participant. During the familiarization procedure, a graphic representing the current incline was presented to the participant. This same graphic was then later used during the choice-selection period to represent the incline on-screen.

After the familiarization period, the choice-selection period began. Eye-movements were recorded using an SR Research Eyelink 1000 Plus sampling at 1000 Hz. Standard 9-point calibration was performed before any data collection, and tracking was routinely verified over the course of the experiment. Choices were always between either one of two referred-to-as reference pairs (4%/6min or 5%/5.5min) and an alternative option of incline and duration varying between 0 to 10% and 0.5 to 10min respectively. 90 total alternative options were presented, each consisting of a different incline and duration as compared to the reference option presented for that trial. After every 90 decisions, subjects were given a short rest of 10 minutes to reduce eye strain. Immediately following this break, a 9-point calibration was again performed to account for any shift in participant posture. Participants were informed that of the selected choice pairs, four were to be randomly selected for the participants to perform after all decisions were made. We further instructed participants that the choice of movement duration held no bearing on the overall time spent for the experimental protocol.

The choice-selection period of the experiment consisted of two distinct phases: the deliberation phase and the decision phase. During the deliberation phase, two graphics representing the choice pairs were presented on-screen at approximately 25% and 75% of the screen width (960-pixel separation, visual angle deviation of ~14°). Participants were given 4.5 seconds to freely deliberate between the two options, and once a preferred choice was in mind, the space bar was pressed to advance to the next phase. If participants had failed to press the spacebar within the allotted timeframe, the trial timed-out and the choice-pairs were repeated at a random later point in the experiment. Furthermore, the participant selection appeared to default to the more metabolically costly of the two, incentivizing participants to make future decisions in time, though these automatic selections were excluded from possible realizations. Should a participant fail to decide within the allotted time, the decision pair was recycled into a random future index for re-selection, and for the given trial the selection defaulted to the more metabolically costly of the two. These automatic selections were excluded from possible realizations.

Next, subjects were required to fixate upon a small cross presented at the center of the screen in order to center their gaze. Participants were required to hold their gaze at this central fixation point for 3 ± 0.25 seconds (mean ± s.d.) before a decision could be made. Fixation duration was randomly selected from a gaussian distribution to discourage anticipatory saccade initiation.

After this fixation duration, the decision phase began. During this phase, two identical grey circles appeared on-screen at the same locations as the previously presented graphics. Participants then were to saccade towards the side of the screen within 1.5 seconds that corresponded to their preferred movement option in the deliberation phase. Failure to initiate a saccade within the allotted time was considered a failure to make a decision, and the trial was repeated at a random later point, and the selection again defaulted to the more metabolically costly of the two. If there was a detected saccade that ended within one of the two circles on-screen, a final screen with the previous corresponding choice pair graphic was displayed to provide feedback and decision confirmation to the participant.

After all choices were made by the subjects, a random sample of four previous choices made were given to participants to then perform. Before doing so, participants waited 15 minutes to account for any variation in time selection between subjects and hold total duration of the protocol constant. Finally, participants walked on the same treadmill previously familiarized on to perform the incline-duration movement pairs randomly selected, again at the same set velocity of 3.2 mph.

### Data processing and statistical analysis

#### Saccade vigor

Eye movement data was converted first from pixel location data to visual angle degrees for both x- and y-coordinates. Instantaneous velocity was calculated via second order Savitzky-Golay filter with a window size of 21 msec. Data was then processed for missing or physiologically impossible saccade velocities. First, samples greater than a threshold of 850 deg/second were identified and the median velocity of all other samples was then calculated. From these high-velocity samples, continuous samples pre- and proceeding were compared to the calculated median value. Each sample greater than the median velocity found within these continuous timeframes were then removed from further consideration. Lastly, all samples with a calculated saccade velocity of zero or *NaN* were also removed from consideration (<1%). Saccades were automatically identified via a post-hoc adaptive velocity threshold algorithm (63). Blinks were identified by continuous samples with missing x- and y- screen coordinate position data. Remaining data was then characterized as fixation periods, where continuous time indices uninterrupted by a saccade or blink taken as indicative of a single fixation event.

As individuals differed in overall saccade velocity and behavior (64, 65), a within-participant measure of saccade vigor was calculated similar to Reppert et. al. (2015). For each participant, the amplitude-velocity relationship was fitted via the following hyperbolic function:

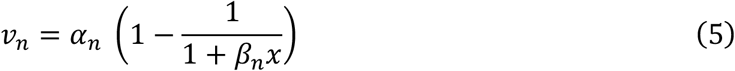

This functional form was chosen as previous work has shown that a hyperbolic relationship is a generally good fit for saccade data (22). Nonlinear Least Squares curve fitting was performed per participant (via *curve_fit* function in the Python *scipy.optimize* package) for both nasal and temporal saccades. Both α and β were free parameters for the curve fit, with *x* the given saccade amplitude, and *v* the saccade peak velocity for participant *n*. Given the fit parameters α and β, the expected peak saccade velocity was calculated. Dividing the measured peak saccade velocity by the expected velocity defined the within-participant vigor metric (referred to as simply *vigor*), with values >1 indicative of more vigorous saccades than typical, and values <1 being less vigorous.

#### Decision making

Participant choices were modelled via general linear mixed models (GLMM) with a logistic link function. Choice response, *SelRef* was coded as a binomial response variable, either 0 for selection of the alternative, or 1 for selection of the reference option; as such, a binomial distribution with logistic link function was used. A typical Gaussian linear mixed-effects model could not be used to ensure that the fitted values ranged from 0 to 1. With the logistic link function, for each choice, the log-odds that participants would select the reference option was estimated. Thus, the model description that for a given choice *i*, participant *j* will choose the reference option is of the form:

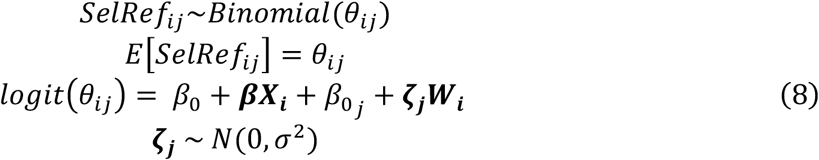

β_0_ is the fixed effect intercept, β the fixed effect coefficients, β_0j_ the participant-specific intercept, ζ_j_ the random effect coefficients, X_i_ the fixed effect parameters for choice *i*, and **W**_i_ the random effect parameters for choice *i*. Regression coefficients and their confidence intervals were estimated via R package *lme4* (66). Random effect coefficients were assumed normally distributed with mean 0 and covariance matrix σ^2^.

The simplified incline/duration mixed logistic model can be reduced to:

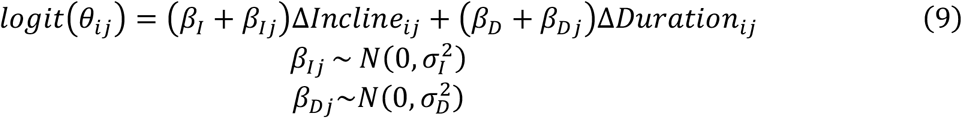

As the ζ parameters above, participants’ individual sensitivities to incline and duration (β_Ij_, β_Dj_) were assumed to be normally distributed across the population with a mean of zero and estimated variances σ_I_^2^ and σ_D_^2^.

To fit the capture rate model of choice preference, a nonlinear mixed effects model was used to fit both participant-specific α values and regression coefficients (Eq 2). Distribution of fitted α values across participants followed a log-normal distribution.

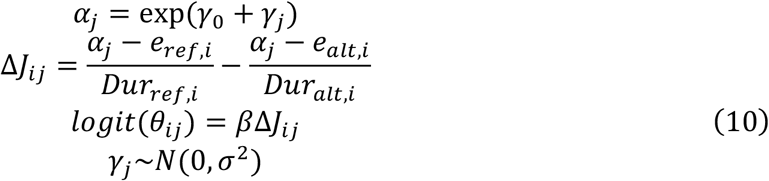

#### Behavioral metrics

Aside from the choice selection, saccade velocity, and other velocity related metrics (e.g. peak velocity, vigor, and trial average vigor acceleration), other measures known to correlate to choice-difficulty, primarily deliberation time and reaction time, were also collected. Participants were provided a maximum of 4.5 seconds from the time of stimulus onset to indicate a choice via keypress. Deliberation time was defined as the time spent from stimulus onset to choice-indication via keypress. If no indication were made, the trial was considered timed out, the choice recycled for a later selection at a random later point in the experiment. Participants were then advanced to the next trial. After successful choice-indication, and the brief fixation period, the decision phase began. Reaction time during this phase was defined as the time from stimulus onset (presentation of two grey circles) to saccade initiation as identified by the algorithm utilized. Should participants time out during the decision phase, the choice was also recycled for later selection. All trials during which participants timed-out either during the deliberation or decision phase were excluded from analysis unless otherwise specified (17.41±20.53 trials; mean±s.d. across participants).

#### Statistical analysis specifications

All statistical analyses were performed using the R programming language, specifically either the *lmer* or *hglm* package for regression analyses. Specifics on regression predictor variables and diagnostics are included in supplementary material. All t-tests performed were two-tailed. Appropriate diagnostics were performed on all regression analysis, ensuring that assumptions regarding homoskedasticity, random effects distributions, and residual distributions were met.

For statistical analysis of deliberation times and reaction times, GLMMs were fitted using Gamma distributed errors with the canonical inverse link function to account for the data’s highly skewed variance structure and to control for heteroskedasticity. Analysis of saccade vigor and rate of vigor change in both phases of the experiment were done via LMM with Gaussian distributed errors.

